# *p*-ClustVal: A Novel *p*-adic Approach for Enhanced Clustering of High-Dimensional Single Cell RNASeq Data

**DOI:** 10.1101/2024.10.18.619153

**Authors:** Parichit Sharma, Sarthak Mishra, Hasan Kurban, Mehmet Dalkilic

**Affiliations:** Luddy Center for Artificial Intelligence, Bloomington, 47408, Indiana, USA; Department of Computer Science, Luddy School of Informatics, Computing & Engineering, Bloomington, 47408, Indiana, USA; Department of Data Science, Luddy School of Informatics, Computing & Engineering, Bloomington, 47408, Indiana, USA; Electrical and Computer Engineering Department, Texas A&M University at Qatar, Doha, 23874, Doha, Qatar; College of Science and Engineering, Hamad Bin Khalifa University, Doha, 23874, Doha, Qatar

**Keywords:** Data-Centric AI, Unsupervised Learning, *p*-Adic Numbers, Clustering High Dimensional Data, Single Cell RNA Sequencing

## Abstract

This paper introduces ***p***-ClustVal, a novel data transformation technique inspired by ***p***-adic number theory that significantly enhances cluster discernibility in genomics data, specifically Single Cell RNA Sequencing (scRNASeq). By leveraging ***p***-adic-valuation, ***p***-ClustVal integrates with and augments widely used clustering algorithms and dimension reduction techniques, amplifying their effectiveness in discovering meaningful structure from data. The transformation uses a data-centric heuristic to determine optimal parameters, without relying on ground truth labels, making it more user-friendly. ***p***-ClustVal reduces overlap between clusters by employing alternate metric spaces inspired by ***p***-adic-valuation, a significant shift from conventional methods. Our comprehensive evaluation spanning 30 experiments and over 1400 observations, shows that ***p***-ClustVal improves performance in 91% of cases, and boosts the performance of classical and state of the art (SOTA) methods. This work contributes to data analytics and genomics by introducing a unique data transformation approach, enhancing downstream clustering algorithms, and providing empirical evidence of ***p***-ClustVal’s efficacy. The study concludes with insights into the limitations of ***p***-ClustVal and future research directions.

## 1 Introduction

Clustering is a pivotal technique that organizes data into meaningful groups, unveiling inherent structures without prior labeling. It is especially critical in real-world applications, like biological data interpretation, where it helps in identifying distinct patterns and groups based on similarities among data points. However, the increasing dimensionality and sparsity of modern datasets pose significant challenges to traditional clustering methods. High-dimensional spaces often dilute the effectiveness of conventional distance metrics and exacerbate the curse of dimensionality, leading to suboptimal clustering performance [1].

In recent times, scRNASeq has marked a paradigm shift, enabling an in-depth exploration of data at an unprecedented single-cell resolution [2]. This technological leap has been instrumental in elucidating complex biological processes, providing vital insights across diverse areas in biology [3, 4, 5, 6]. A primary challenge in scRNASeq data analysis is identifying distinct cell groups, akin to unsupervised clustering in machine learning [7]. The field has seen diverse clustering methodologies evolve, each with its own unique approach and data processing strategy [8, 9].

Despite its potential, scRNASeq data processing is beset with challenges, primarily due to the noisy, low-resolution nature of the raw data, typically represented as an *m ×d* matrix of cells (samples) and gene counts (features). For instance, gene expression is often captured across thousands of genes (dimensions), clustering such high dimensional space is difficult due to distance measures becoming less meaningful in distinguishing data points [1]. scRANSeq data is characteristically sparse i.e. several genes have zero/near-zero counts across the cells, thus making it difficult to find meaningful structures, or establish similarity between data points [3]. Moreover, noise is introduced in the form of dropout events (when a gene expression is not captured even when it’s present), due to technical limitation and biological variation [8]. This makes it difficult for clustering algorithms to separate true gene expression from nuisance variables. Due to biological variability (cell development stages, cell cycle variability etc.), there is sufficient variation within cells that can blur boundaries between cell types and states, making clustering more challenging [2]. Consequently, existing clustering algorithms often struggle to effectively cluster such complex data spaces. For completeness, we share a general overview of the standard data acquisition and processing pipeline in Figure 1.

**Fig. 1:**
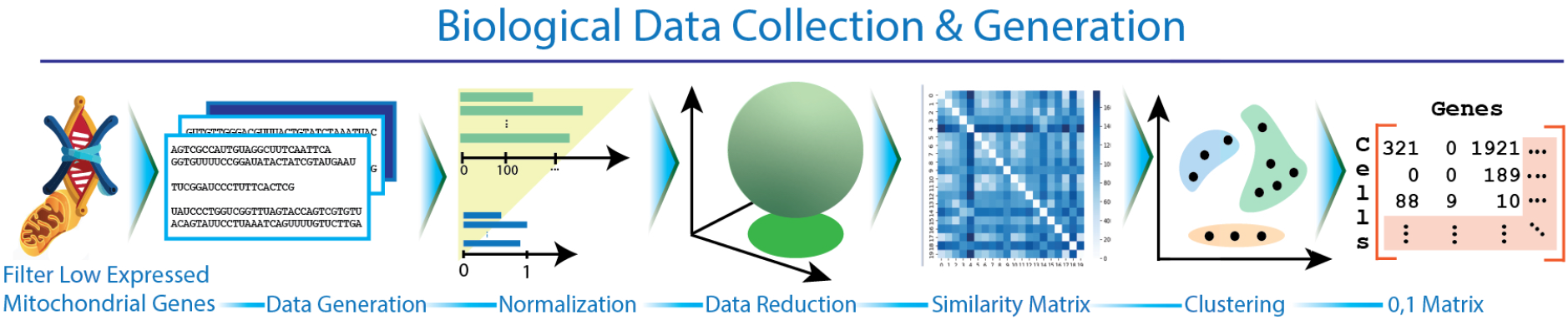
Data generation phase. The samples are extracted from the tissue of interest, followed by cell library preparation and high throughput sequencing. The obtained sequences are aligned with a reference genome. The result is a matrix that stores gene transcripts counts per cell. **Data analysis phase**: The data goes through initial quality control to remove low quality cells and genes, normalization, feature subset selection or dimensionality reduction, building cell similarity matrix either in the lower dimensional or original feature space, and clustering.

In this work, we introduce *p*-ClustVal, a novel data transformation technique inspired by the *p*-adic number theory [10]. By leveraging alternate metric spaces based on *p*-adic-valuation, *p*-ClustVal enhances cluster discernibility and separation, setting it apart from conventional methods. By operating in the transformed *p*-adic space, *p*-ClustVal mitigates cluster overlap, thereby enhancing cluster separation. This transformation consequently augments the efficacy of downstream clustering algorithms, facilitating the elucidation of previously obscured patterns and structures within the data. One of the key strengths of *p*-ClustVal lies in its ability to seamlessly integrate with and enhance popular clustering algorithms and dimension reduction techniques. Moreover, *p*-ClustVal is designed to be user-friendly, requiring minimal parameter tuning. It employs a unsupervised heuristic for dynamically selecting optimal parameters without requiring ground truth labels, making it more accessible. To validate the effectiveness of *p*-ClustVal in real-world applications, we apply it on several high-dimensional, and sparse datasets in scRNASeq datasets.

Extensive evaluation across 30 experiments comprising 1450 observations reveal that *p*-ClustVal significantly improves the performance in 91% of cases, with the clustering labels showing a much higher level of agreement with the ground truth. The primary contributions of this work include:

- A novel transformation (*p*-ClustVal) designed to improve cluster identification in high dimensional single cell data.
- Introducing a unsupervised heuristic to find optimal transformation parameters in a data-driven manner, significantly improving adaptability across datasets and reducing manual configuration.
- An intuitive demonstration of *p*-ClustVal on synthetic and real world data, and discussion of essential metric properties.
- Comprehensive empirical validation of *p*-ClustVal’s efficacy in enhancing classical and state of the art clustering tools.

The remainder of this paper is organized as follows: Section 2 outlines the background and related work. Section 3 elaborates on our proposed method. Section 4 details the experimental setup, and Section 5 presents our findings. Section 6 addresses the current limitations and explores potential solutions. Finally, Section 7 concludes the paper, offering final thoughts and directions for future research.

## 2 Related Work

Contemporary analysis of scRNA-seq data encompasses diverse computational stratification methodologies, broadly categorized into: partitional algorithms enforcing strict boundaries, hierarchical approaches constructing nested groupings, graph-theoretic frameworks with community detection, and deep neural architectures. Single-cell datasets inherently harbor stochasticity from both technical artifacts (sequencing bias, library preparation) and biological variation [11]. To deconvolve biological signals from technical noise, dimensionality reduction is employed prior to clustering, simultaneously attenuating the curse of dimensionality and computational complexity. Here, we present a systematic analysis of prevalent dimensionality reduction approaches and clustering paradigms, examining their algorithmic foundations and limitations.

### Dimensionality reduction techniques

Prevalent dimensionality reduction approaches—including Principal Component Analysis (PCA), T-Distributed Stochastic Neighbor Embedding (t-SNE), and Uniform Manifold Approximation and Projection (UMAP). These can function both independently and in conjunction with clustering algorithms to mitigate noise, enhance clustering efficacy, and optimize computational efficiency. Deep neural autoencoders have recently emerged as an alternative paradigm for simultaneous dimensionality reduction and clustering, discussed subsequently in clustering methodologies.

- **PCA** [12] implements orthogonal transformation to project high-dimensional data onto a lower-dimensional subspace defined by principal components, which are eigenvectors maximizing explained variance. Dimensionality reduction is achieved through selective retention of dominant eigenvectors; however, the method’s underlying assumption of linear correlations renders it susceptible to missing intrinsic non-linear manifold structures [13].
- **t-SNE** [14] employs non-linear manifold learning to project high-dimensional data onto lower-dimensional embeddings while preserving topological structure. The algorithm optimizes a probabilistic framework by minimizing the Kullback-Leibler divergence between joint probability distributions computed in both high and low-dimensional spaces. While successfully implemented in single-cell transcriptomics analyses [15, 16], t-SNE exhibits notable limitations: quadratic computational complexity *O*(*n*^2^) impeding scalability, stochastic initialization leading to non-deterministic outputs, and potential loss of global topological relationships during dimensionality reduction.
- **UMAP** [17] implements a manifold learning framework predicated on algebraic topology and Riemannian geometry to preserve both local and global data structures during dimensionality reduction. The algorithm constructs weighted simplicial complexes in the original space, followed by optimization of a fuzzy topological representation that captures multi-scale relationships. Through stochastic gradient descent, it minimizes the cross-entropy between high and low-dimensional topological embeddings. While demonstrating superior performance to t-SNE [18] and exhibiting enhanced scalability, UMAP’s efficacy is contingent upon hyperparameter optimization, and its stochastic nature introduces non-deterministic outcomes across iterations.

### Partition based clustering

Several algorithmic variations enhance k-means’ capabilities: pcaReduce [19] employs hierarchical merging of clusters on principal components while dynamically optimizing cluster numbers through iterative PC removal; SAIC [20] performs feature selection and automatic cluster number adjustment; SCUBA [21] extends k-means to temporal analysis by incorporating bifurcation detection in developmental trajectories. To address the inefficacy of Euclidean metrics in high dimensions [1], RaceID3 [22] implements k-medoids [23] with Pearson correlation distances.

### Hierarchical clustering

Hierarchical clustering methodologies encompass both agglomerative (bottom-up) and divisive (top-down) approaches to generate dendrogram-structured data partitions [24]. Contemporary computational implementations augment these foundational methods: BackSpin [25] employs PCA-based dimensionality reduction followed by iterative binary partitioning via modified k-means; CIDR [26] implements dropout imputation and principal coordinate analysis to address data sparsity and high dimensionality while mitigating outlier sensitivity; SC3 synthesizes hierarchical clustering with consensus matrices to optimize k-means-derived partitions.

### Graph & community detection methods

Graph-theoretic clustering approaches leverage topological data representations through nearest-neighbor connectivity to identify community structures. The Louvian algorithm [27] facilitates parameter-free community detection, prominently implemented in Seurat [28], which constructs refined shared nearest-neighbor graphs from PCA-reduced expression data before community partitioning. Spectral clustering [29] transforms data into an eigenspace via graph Laplacian computation to enhance cluster delineation, while SIMLR employs multiple kernel metrics to learn cell-cell similarity measures, thereby achieving robustness to noise and heterogeneity while transcending predefined metric limitations.

### Deep learning approaches

architectures have emerged as powerful frameworks for single-cell clustering. scDeepCluster integrates autoencoder networks with dedicated clustering layers, implementing self-training mechanisms via pseudo-label optimization to address dimensionality reduction and dropout compensation simultaneously. scziDesk [32] incorporates zero-inflated negative binomial distribution within an autoencoder framework, augmented by discriminative loss functions, to model dropout events while learning latent representations. ADClust [33] employs pre-trained autoen-coders for dimensionality reduction followed by statistical merging of micro-clusters, enabling automated determination of optimal cluster cardinality without a priori specification.

### Miscellaneous

Weighted ensemble methods attempt to improve cell heterogeneity analysis by integrating multiple clustering results, often weighted by the reliability of base clustering. SAFE [34] combines four state-of-the-art clustering techniques (SC3, CIDR, Seurat, t-SNE + *k*-means), and use hyper-graph partitioning to ensemble the results into a consensus matrix. sc-GPE [35] combines graph based clustering techniques to construct the consensus matrix, followed by hierarchical clustering on consensus matrix. scEWE [36] proposed higher-order element-wise weighting strategy, to assign different clustering weights to different cells, leading to a robust consensus matrix. scEWE mitigates the issue of ignoring low-level structure information (cell-cell relationships) in previous ensemble methods by assigning different weights to cells.

jSRC [37] perform joint optimization of dimension reduction and clustering to address scalability and clustering accuracy. [38] use LDA (latent Dirichilet Allocation) to identify genes representative of various clusters, thus enabling data-driven cluster annotation. SCENA [39] select specific genes (feature set), followed by improving local affinity amongst cells and clustering based on a consensus matrix. FEATS [40] focus on selecting top informative features for downstream clustering, cluster number annotation and outlier detection. scHFC [41] combine Fuzzy C-means (FCM) and Gath-Geva (GG) algorithms with simulated annealing and genetic algorithm for clustering the scRNASeq data.

### Application of p-adic Theory

*p*-adic number theory, fundamental to algebraic arithmetic, has demonstrated emergent applications across diverse computational and biological domains. Recent investigations have leveraged p-adic spatial hierarchies for network topology interpretation [42], particularly in Erdős-Rényi graph theoretic simulations. Furthermore, p-adic metrics have elucidated genetic code structures through analysis of codon proximity via 5-adic and 2-adic distances [43], while contemporary research has established connections between p-adic statistical field theories and neural architectures [44, 45].

Despite these advances, the integration of *p*-adic numbers in machine learning is still nascent. In this study, we draw inspiration from the work of [46] on *p*-adic vectors and propose an innovative approach to enhance unsupervised clustering. Previous efforts have been made to use the natural hierarchical properties of *p*-adic numbers for sparse encoding, thus improving clustering [47, 48]. Some authors have also used the *p*-adic metric instead of Euclidean, Manhattan, Chebyshev, or cosine metrics to improve k-nearest neighbor [49]. Here, we present a different transformation inspired by the p-adic valuation. Our method is able to discretize the space using parameters inferred from the data, thus introducing a novel data representation.

### Comparison with existing transformation methods

As discussed earlier, data transformation methods come with their unique strengths and drawbacks. For example, assumption of linearity (PCA) versus non-linearity (t-SNE/UMAP) affects how well a given method can discover complex non-linear relationships [50]. Preservation of global (PCA/UMAP) versus local structures (t-SNE) determines if the results would conserve overall structure or emphasize local fine-grained relationships [18]. Choice of dimensionality reduction method also affect the technical noise and computational cost differently [51]. Since, many clustering tools include dimensionality reduction (PCA, PCoA, deep neural autoencoders etc.), the results are also affected by the choice of algorithm [52]. In contrast, *p*-ClustVal does not assume linearity/non-linearity, rather than explicitly preserving global vs. local structure, the emphasis is to increase cluster separability in the transformed space. Experiments show that, where performance of dimensionality reduction methods may vary with data, *p*-ClustVal improves clustering accuracy across data and dimensionality reduction (Sec. 5). This indicates that *p*-ClustVal is robust to data specific characterstics. Unlike some methods where final results vary due to stochastic nature of the algorithm (t-SNE/UMAP), *p*-ClustVal follows deterministic process that always yield same results. Moreover, compared to prevalent dimensionality reduction techniques, *p*-ClustVal is computationally efficient, as it scales well with data size, and relatively cheap to compute. Further details on complexity analysis are given in Sec. 3.7.

## 3 Methods

We define the notation as follows: let **x ∈**ℝ^*d*^ represent a *d*-dimensional vector over the reals. The dataset *D* = {**x**_1_, **x**_2_, …, **x**_*m*_} comprises *m* such vectors. We denote the *i*^th^ element of **x** by *x*_*i*_. The distance between two vectors **x** and **y** is given by *d*(**x, y**) = ∥**x**− **y**∥ _2_. The number of centroids is denoted by *k*∈ ℕ, and we also use *k* to denote the top *k* neighbors of a data point, with the specific usage clarified by the context.

### 3.1 *p*-adic numbers

*P*-adic numbers were first introduced by Kurt Hensel in 1897. *P* -adic numbers are an extension of rational numbers obtained by completing the rational numbers with respect to the *p*-adic metric. They are represented as infinite series *a*_0_ + *a*_1_*p* + *a*_2_*p*^2^ + …, where *a*_*i*_ are integers and *p* is a prime number. Convergence of the series is determined by the size of the powers of *p*. A *p*-adic series will converge when the terms become arbitrarily small as the index increases, measured by the *p*-adic-valuation. *p*-adic-valuation (*V*_*p*_(*n*)) [10, 53] is a mathematical function that finds the highest power of a prime number *p* that divides a given non-zero rational number *n*. It is defined as follows:

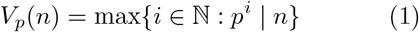

Given the diverse applications of *p*-adic numbers in scientific domains, and considering the capability of *p*-adic-valuation to provide alternative data representations with respect to a prime *p*, we introduce a parametric space discretization technique inspired by *p*-adic-valuation. Specifically, our transformation function discretizes data to enhance the separation between different classes while preserving the proximity of data points within the same class.

### 3.2 *p*-ClustVal: Addressing the Limitations of *p*-adic-valuation

In *p*-adic-valuation, the largest exponent of a prime that divides a number is identified. For instance, the 2-adic valuation of 3 is 0, because the highest power of 2 that divides 3 is 0 (2^0^ = 1). Consider a set of data points, *X* = {3, 5, 11, 67, 129, 259}. Calculating the 2-adic valuation results in *V*_2_(*X*) = {0, 0, 0, 0, 0, 0}, *i.e*., all data points are mapped to 0, losing the inherent distinctions in the data. Such loss of distinction compromises clustering; for instance, if *k* = 2 and *k*-means is done on *V*_2_(*X*), it would erroneously group all points into a single cluster. This is an undesirable outcome, because after the valuation, we would ideally want data points 3, 5, 11 to be assigned a different cluster than 67, 129, 259. On the contrary, *p*-adic-valuation in it’s current form would force any distance based clustering to assign these points as one cluster. Therefore, to transform a given number *x* (while preserving relative order of magnitude), we find the highest power of a reference number such that reference raised to that power is smaller than the given number *x*. We relax the constraint of using prime number, thus allowing the reference to be any real number, and refer to it as cohesion (`coh`). We call this new transformation as *p*-ClustVal. Using this new adaptation and `coh` set to 2, *p*-ClustVal({*X* = 3, 5, 11, 67, 129, 259}) → {1, 2, 3, 6, 7, 8}, reflecting that 2^1^ is the highest power of 2 which is less than 3, 2^2^ is less than 5, 2^3^ is less than 11, 2^6^ is less than 67, and similarly for other values in *X*.

With this new valuation, numbers 1, 2, 3 are closer to each other, similarly 6, 7, 8 have become closer. If *k*-means is run on this new valuation, naturally it leads to better clustering because numbers 1, 2, 3 are further from numbers 6, 7, 8. Since all the numbers between *coh*^*n*^ to *coh*^*n*+1^ will end up at *n*, our transformation preserves relative monotonicity of datum’s magnitude. This enhances the clustering accuracy since the transformed values corresponds better with the magnitude of data.

### 3.3 Transformation Function

We define a transformation function *θ* : ℝ →*Z*, parameterized by separation (`sep`) and cohesion (`coh`), for any scalar *x*_*i*_ ∈ℝ. For simplicity, here we denote `coh` by *c*, and `sep` by *s*. This function is given by:

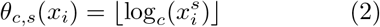

The transformation is applied element wise to every *x*_*i*_ of vector **x**, such that *s* scales each dimension (*d*) of **x**, spreading the data points apart, while *c* maps the scaled data to *c*^*d*^ hypercubes. After the transformation, we further separate the data via uniform expansion (*θ*_*c,s*_(**x**)^2^). Since the transformation is done element wise, the run time complexity is linear in the size of data i.e. *O*(*md*).

### 3.4 Preservation of Distance Metric

Our transformation *θ* is designed to preserve the properties of the metric space:

#### Identity of Indiscernibles

For a datum **x** in the input space, we have *d*(**x, x**) = 0. This property is preserved post-transformation:

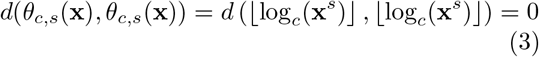

#### Positivity

For datum **x, y**, with *d*(**x, y**) *>* 0, their transformed counterparts also maintain a positive distance:

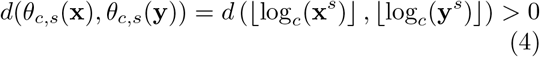

This holds unless **x** and **y** are mapped to the same point by *θ*. If **x** and **y** are mapped to the same point, then distance between them is 0 by Eq. (3). **Symmetry:** For datum **x, y** in the input space, satisfying *d*(**x, y**) = *d*(**y, x**), the transformed space retains this symmetry:

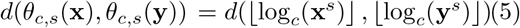

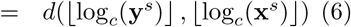

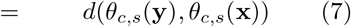

The symmetry is maintained as the transformation is consistent for all points.

#### Triangle Inequality

For **x, y** and z, if *d* (x, y) + *d* (y, z) ≥ *d* (x, z), then:

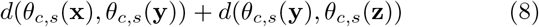

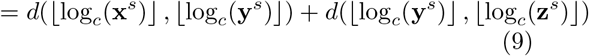

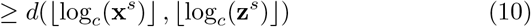

This is proven by the monotonic nature of the transformation. Let **n** = *θ*_*c,s*_(**x**), **m** = *θ*_*c,s*_(**y**), **z** = *θ*_*c,s*_(**z**). If *d*(**0, y**) *> d*(**0, x**), then *d*(**0**, └log^*c*^(**y**^*s*^)┘) *> d*(**0**, └log^*c*^(**x**^*s*^)┘). Similarly, for **z** and **y**, and **z** and **x**. Therefore, for transformed points **m, n**, and **k**, *d*(**m, n**) + *d*(**m, k**) ≥ *d*(**n, k**), maintaining the triangle inequality in the transformed space.

### 3.5 Illustrative examples

Consider the example in Fig. 2-(A), which shows data in two dimensions with points in violet and orange representing two different clusters. Points `G`, `H`, and `I` are equidistant from members of both clusters. Moreover, `H` is closer to `C1` than `C2`, and if the points are assigned based on their proximity to the centers then `H` will be misclassified as part of `C1`. Fig. 2-(B) illustrates the data after scaling by `sep` = 3. Though, scaling pushes data points apart, it does not reduce the misclassification risk, because the spatial orientation of data is unaffected. This issue is addressed via discretization with `coh` = 2 (Fig. 2-(C)). The transformation relocates boundary points `G`, `H`, and `I` closer to their actual center, significantly enhancing the clustering accuracy. Data points are mapped to the edges of squares of dimension `coh` x `coh`, and some points, for instance: `M, N` are mapped to the same location in the transformed space. This is because for each dimension in the actual data, the transformation process is essentially collapsing all points between *coh*^*n*^ to *coh*^*n*+1^ at *n*. Although, the example is illustrated in two dimensions, same process is extensible to higher dimensions by applying it on each dimension individually.

**Fig. 2:**
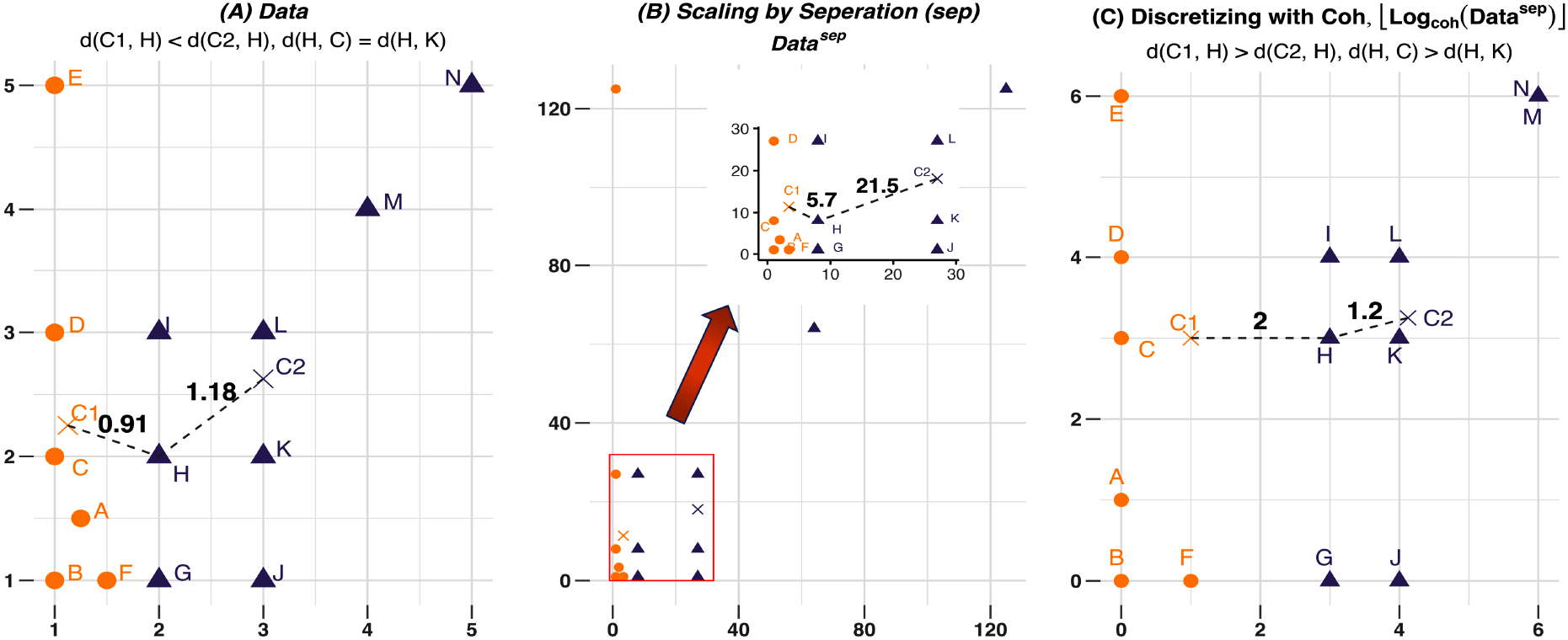
Visualizing the affect of transformation on a simulated example. Orange and Violet points represent two different clusters. *C*1 and *C*2 are the respective centroids. (A): actual data, (B): after scaling the data with `sep` = 3. Figure in the inset highlights the data within the red region. (C): after discretizing with `coh` = 2. Dashed lines indicate the distance between cluster centers *C*1, *C*2 and *H*. Scaling pushes data points apart, but conserves the global spatial structure of data, as seen in plot-(B). Discretizing with `coh` brings data points in the same cluster closer, thus increasing the separation between different clusters.

The example intuitively explains the advantage of using *p*-ClustVal in conjunction with clustering. However, the application is not limited to simulated data, and we show that similar behaviour can be obtained on real world data.

Fig. 3 visualizes the affect of transforming the Muraro dataset (details given in Table 1). The data has 9 classes as shown by the types in the legend. The points are colored by true cluster labels and the circles represent the 85% confidence interval for the data. The first plot (top) shows the actual data, and the second plot (bottom) shows the transformed data on the top two principal components. The overlap between clusters, for instance: `PP cell and A cell`, is noticeably reduced in transformed data. Additionally, for clusters where the spread of the data was high, for example: `Acinar cell, Ductal cell`, the spread is visibly reduced and clusters have become significantly compact after the transformation. Specifically, data points in cluster `PP cell` have gotten closer while moving away from neighboring clusters. Spread of points in `Stellate cell` has also reduced. Compared to the actual data, the transformation has noticeably improved the chances of clustering `A cell, PP cell, Stellate cell, Acinar cell` and `Ductal cell`. The visual interpretation is also supported by the quantitative assessment on several real world datasets (Sec. 4).

**Table 1:** Details of scRNASeq datasets (post processing) including size, features, and cluster count.

| Data | $m$ | $d$ | $k$ | Sparsity | Source |
| --- | --- | --- | --- | --- | --- |
| Pollen | 301 | 19952 | 11 | 72.72 | [60] |
| Darmanis | 420 | 19422 | 8 | 78.89 | [61] |
| Usoskin | 622 | 18023 | 4 | 80.05 | [62] |
| Mouse Pan-creatic | 1884 | 11900 | 13 | 86.31 | [63] |
| Muraro | 2122 | 15323 | 9 | 66.52 | [64] |
| Limb | 1090 | 16624 | 6 | 85.22 | [65] |
| Trachea | 1350 | 18347 | 4 | 81.54 | [65] |
| Lung | 1676 | 17243 | 11 | 85.23 | [65] |
| Diaphragm | 870 | 15813 | 5 | 87.26 | [65] |
| Spleen | 9552 | 9705 | 5 | 86.50 | [65] |

‘ $m$ ’: number of data points (or cells), ‘ $d$ ’: number of dimensions (or genes), $k$ : true cluster count. Sparsity indicate percentage of zero values in data.

**Fig. 3:**
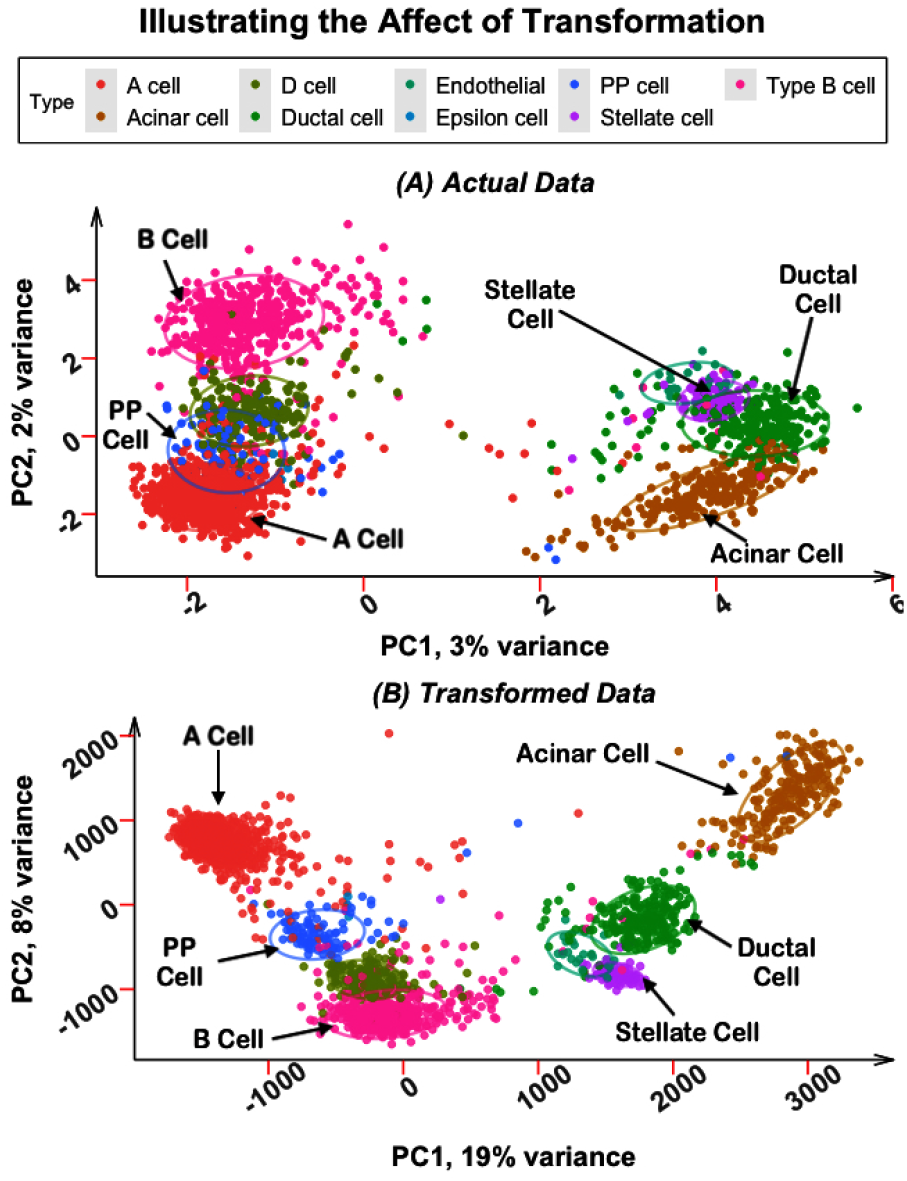
Visualization of Muraro data on first two principal components. (A): actual data, (B): transformed data. Post transformation, overlap between different clusters has lessened driven by the reduction in spread of data within clusters, and separation between clusters have increased.

### 3.6 Data-centric Search for Optimal Parameters

*p*-ClustVal effectuates n-dimensional hypercubic manifolds of volume `coh`^*n*^, mapping datapoints to vertices; however, optimal `coh` parameterization without ground truth labels presents a fundamental optimization challenge. Excessive hypercubic volume risks collapsing distinct modalities through over-compression, while insufficient volume fails to aggregate proximate points, potentially leading to inter-cluster overlap.

In the presence of ground truth labels, classspecific `coh` values can be determined by setting `coh` to the maximum intra-class spread, ensuring all points within a class are mapped to their respective bins and improving clustering accuracy. However, real-world data often lack labels, necessitating an unsupervised heuristic to dynamically determine parameters from the data. The objective is to enable *p*-ClustVal to automatically adapt the parameter search to different datasets while minimizing manual tuning. As `coh` is influenced by the overall data spread, an automated method to probe the dataset is required.

Setting `coh` to the average distance between a data point and its *k* nearest neighbors provides a global estimate of proximity within the entire dataset, as defined in (11). The choice of *k* affects the neighborhood and, consequently, the estimate of `coh`; small *k* values may result in insufficient points being mapped to the same hypercube, while large *k* values may inflate *coh* and collapse points from neighboring clusters. The number of neighbors was selected empirically by repeating the clustering on different *k* values for 5 randomly selected datasets, with *k* = 5 chosen for the experiments based on the similar range of ARI obtained (Appendix A).

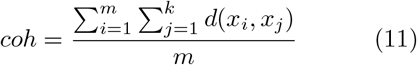

*k* denote the number of neighbors and *x*_*i*_ ≠ *x*_*j*_.

Overly large features in single cell data [54] can lead to large distances between closest points, inflating the estimate of `coh` and causing non-numeric results during discretization when using *knn* search on raw data. To address this issue, *knn* search is performed on standardized data, preserving the original neighborhood [55] and avoiding excessively large `coh` values. The separation parameter (`sep`) is set to `coh`+1 to expand the space by at least `coh`, pushing points that do not belong to the same cluster further apart. Since `coh` *>* 0 and `sep` *>* `coh`, raising data to a positive exponent higher than 1 always expands the data space. Figure 4 illustrates the steps involved in the algorithmic workflow.

**Fig. 4:**
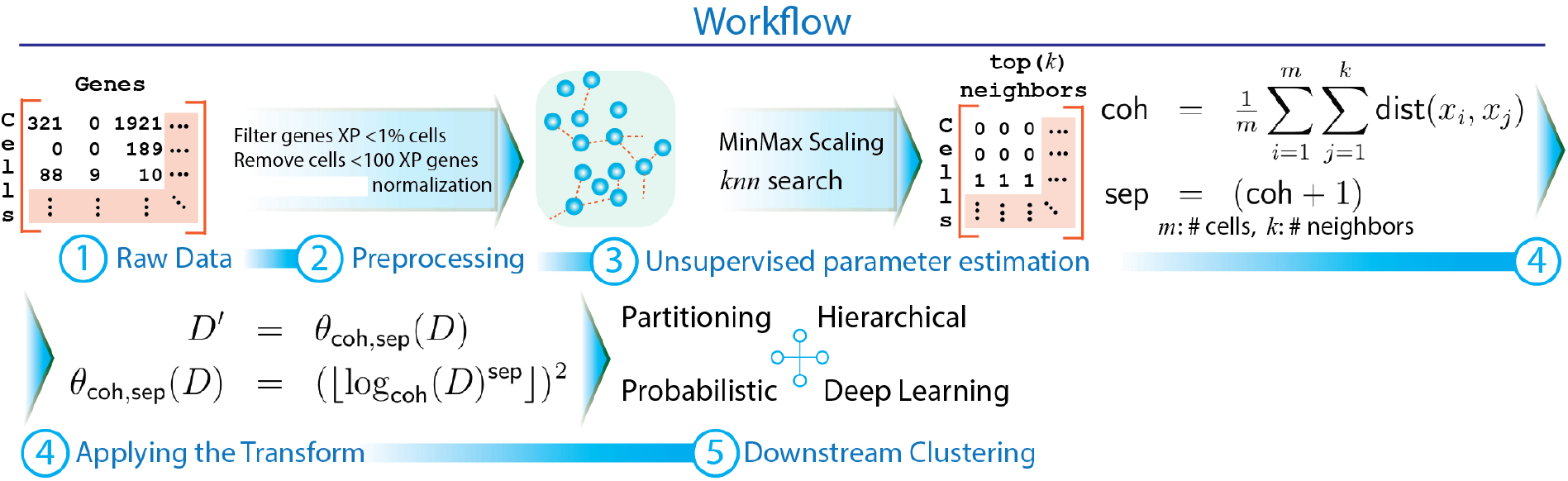
Algorithmic workflow in *p*-ClustVal: (Steps 1-3), processing the raw data to filter low quality cells and genes, followed by data normalization and scaling. (Step 4), finding the optimal parameters in a data-centric manner and applying the *p*-ClustVal transform. (Step 5), applying clustering on the transformed data.

### 3.2 Complexity Analysis

*p*-ClustVal is an element wise transformation, meaning that it’s time complexity is linear in the size of data (*O*(*md*) where, *m* : number of samples and *d* : number of dimensions). However, the total time is also affected by the parameter search process. The time complexity of parameter search is dominated by the neighbor search (*O*(*m*^2^*d*)). We obtain the time complexity of our approach by adding the component parts: *O*(*md*) + *O*(*m*^2^*d*) ≈ *O*(*m*^2^*d*). Though, total complexity is quadratic, yet *p*-ClustVal is practically efficient because parameter search is one time process. We benchmarked the time usage of *p*-ClustVal on large synthetic data [56], and results are shown in Fig. 5. Red line indicate the average time (average of 5 trials) required for *p*-ClustVal. Across datasets, runtime of *p*-ClustVal is well below 1 minute. In-fact, on the largest dataset with 100, 000 points, *p*-ClustVal only takes 57 seconds. The cyan line show the average total time required for executing kNN neighbor search, estimating optimal parameter values and performing the *p*-ClustVal transformation. On the largest dataset (with 100K points), the total time is 12 minutes. Although, neighbor searches can be executed more efficiently [57, 58, 59], in it’s current form, *p*-ClustVal only takes few minutes to complete the process. We defer the exploration of faster neighbor search to future work.

**Fig. 5:**
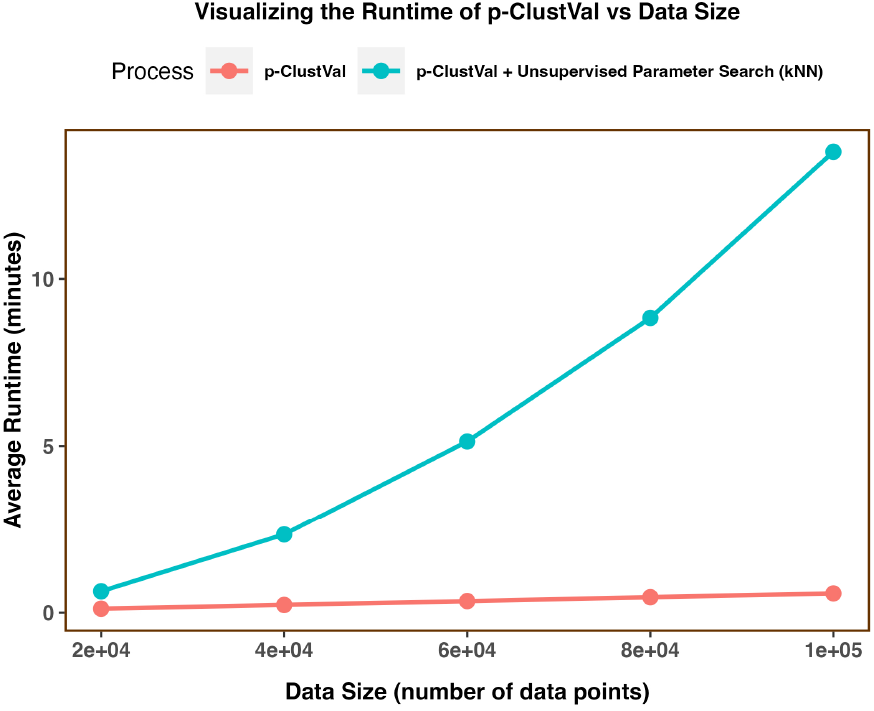
Scalability analysis on larger datasets. *p*-ClustVal transform finishes within a minute on all datasets. Although, parameter search is relatively time consuming, but it’s a one time process. Total time for neighbor search, parameter estimation and *p*-ClustVal is few minutes across datasets.

## 4 Experiments

Experiments on 10 high-dimensional and sparse scRNASeq datasets (Table 1) demonstrate the improvement in clustering accuracy by utilizing transformed versus raw data. Additionally, we show that the efficacy of widely used dimension reduction techniques and the quality of results in SOTA scRNASeq clustering are enhanced by applying the transformation. As described in Sec. 2, SOTA tools differ greatly in terms of the underlying clustering and data processing. To minimize the impact of different data processing pipelines, the first two experiments use the same data processing and focus on directly comparing the clustering algorithms forming the backbone of existing scRNASeq methods, while the third experiment compares domain-specific tools using their default data processing (Appendix C and D).

### 4.1 Benchmark 1: Enhancing Classical Clustering

The experiments aim to determine if *p*-ClustVal can improve the accuracy of classical clustering methods, demonstrating that data transformed through *p*-ClustVal and clustered using a simple algorithm like *k*-means can outperform more sophisticated algorithms applied to raw data. The second part of the experiment shows that *p*-ClustVal enhances the performance of most algorithms, confirming its compatibility and complementarity with widely used clustering techniques. Representative algorithms from partition-based clustering (*k*-means, Fuzzy *k*-means (FZKM)), mixture model-based clustering (expectation-maximization algorithm (EM)), and hierarchical clustering (ward agglomerative clustering (WAC)) are selected. SOTA implementations of other techniques are compared in Sec.,5. Results are evaluated over 20 trials and true cluster count for each dataset, with all algorithms initialized using the same centroids (*k*++ initialization) in a given trial. Box-plots in Fig.,6 show summary statistics, and average statistics are reported in Table,2.

### 4.2 Benchmark 2: Clustering with Dimension Reduction

Dimension reduction, a common pre-processing step to filter noise or obtain a lower-dimensional representation of data before clustering [4, 26, 66]. Here, we show that performance is improved by using transformed data. Three frequently used algorithms, namely Principal Component Analysis (PCA), t-distributed Stochastic Neighbor Embedding (t-SNE) [14], and Uniform Manifold Approximation and Projection (UMAP) [17], are compared. Results demonstrate that applying dimension reduction on transformed data greatly improves clustering accuracy in the majority of datasets (Sec.5). The number of components is selected to capture 90% variance in data for PCA, while t-SNE and UMAP components are fixed at 2. After applying dimension reduction to raw and transformed data, 20 trials of *k*-means clustering are performed, keeping centroids (*k*++ initialization) the same in a given trial, and average statistics are reported.

### 4.3 Benchmark 3: Enhancing the state-of-the-art in Domain Specific Clustering

Due to the large number of available tools for clustering single cell data, including packages based on classical clustering or its variants, packages that perform clustering after projecting data to lower-dimensional space, and approaches based on deep learning models to extract latent embeddings for clustering, an exhaustive comparison is difficult. Therefore, evaluation is done via representative approaches in each category. Popular packages compared include Seurat [28], which uses network-based community detection; RaceID (*k*-Medoids clustering) [22]; SIMLR [67], which uses spectral clustering; SC3 [68], which combines dimension reduction with multiple matrices (Euclidean, Pearson coefficient, etc.); and ADClust [33], which performs clustering on low-dimensional representations obtained from auto-encoding deep neural networks.

### 4.4 Evaluation criteria

Quality of clustering assignments is evaluated via Adjusted Rand Index (ARI). Specifically, clustering assignments are compared with ground truth, and ARI is reported. ARI measure the extant of similarity between different clustering assignments and it’s value range from ™0.5 to 1, where 1 indicates identical clustering, 0 represents random clustering, and negative values indicate less agreement than expected by chance.

## 5 Results & Discussion

### p-ClustVal improves k-means

Results (Fig. 6) show that *k*-means on transformed data (T-*k*-means) performs significantly better than other methods. Additionally, T-*k*-means achieves noticeable gains in accuracy on specific datasets, e.g. Usokin (T-*k*-means: 0.45, second best (fz-*k*-means): 0.10), Limb (T-*k*-means: 0.66, second best (*k*-means): 0.14), Trachea (T-*k*-means 0.58, second best (WAC): 0.23), Lung (T-*k*-means: 0.49, second best (WAC): 0.19) and Diaphragm (T-*k*-means: 0.83, second best (*k*-means): 0.25). Results demonstrate that *p*-ClustVal can greatly boost the clustering accuracy of *k*-means. Interestingly, the observed gain is conserved across datasets, which further underscores the robustness of *p*-ClustVal to varying data sizes, number of dimensions and number of clusters.

**Fig. 6:**
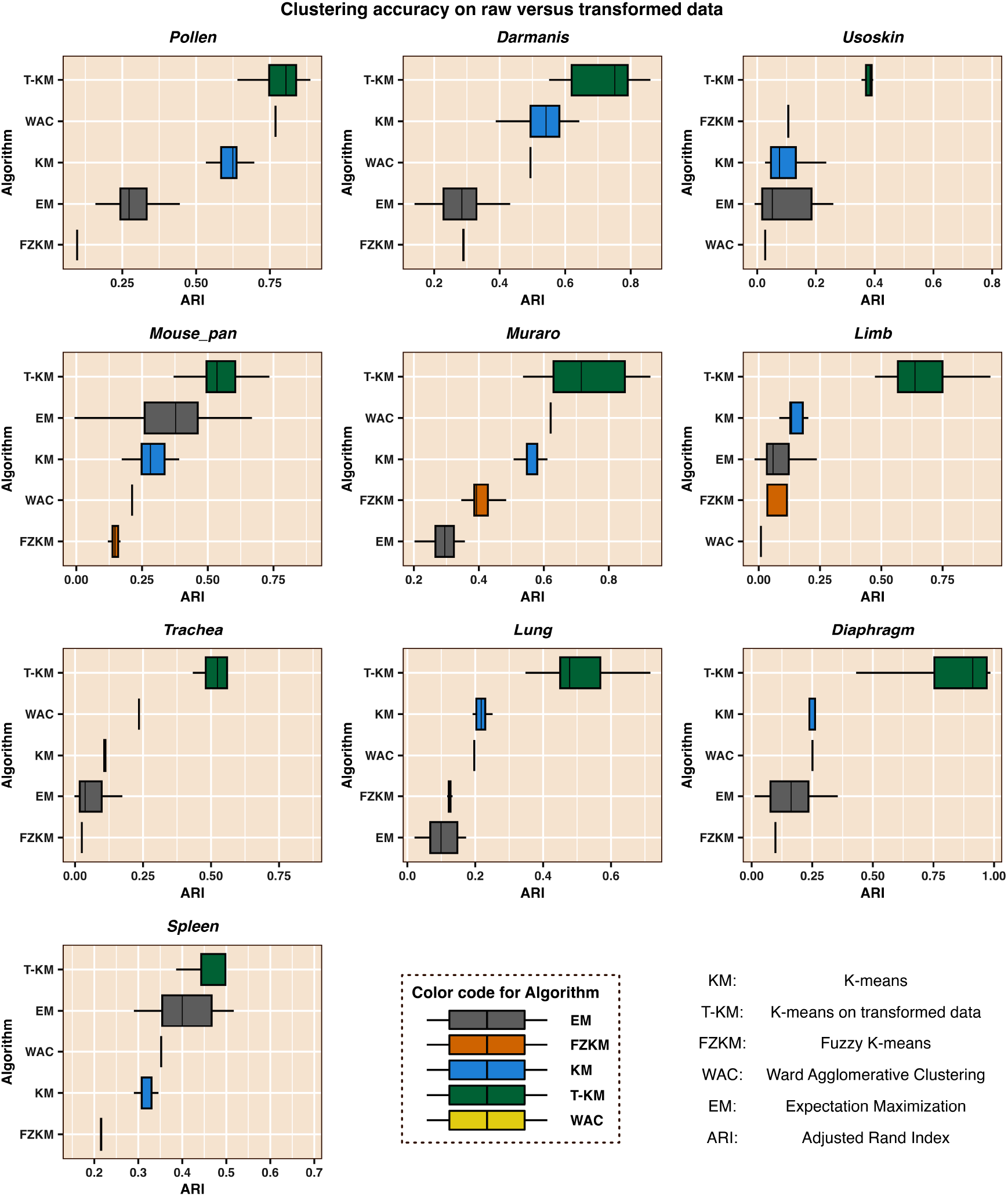
T-*k*-means augmented with *p*-ClustVal performs significantly better, and surpass other algorithms. The gain in accuracy is consistent across datasets and *k*.

### p-ClustVal enhances classical clustering

The results presented in Table 2 underscore the importance of *p*-ClustVal in enhancing clustering performance. While EM on raw data outperformed its transformed counterpart on the Spleen dataset, WAC applied to the transformed data achieved substantially higher ARI scores, surpassing EM by a notable margin of 0.36. Across datasets, the transformation’s impact is further highlighted by the maximum ARI improvements observed, with *k*-means’s ARI increasing by 0.58 on the Diaphragm dataset, Fuzzy-*k*-means exhibiting a remarkable 0.67 improvement on the Limb dataset, EM’s ARI increasing by 0.46 and 0.48 on both the Usokin and Diaphragm datasets, and WAC’s performance being enhanced by 0.72 on the Diaphragm dataset. The consistent improvement in ARI scores of clustering algorithms on the majority of datasets demonstrates the effectiveness of *p*-ClustVal.

**Table 2:** Performance of different clustering techniques on raw versus transformed data.

| Dataset | Adjusted Rand Index (ARI) ± S.D. | | | | | | | |
| --- | --- | --- | --- | --- | --- | --- | --- | --- |
|  | *k*-means |  | FZKM |  | EM |  | WAC |  |
|  | *R* | *T* | *R* | *T* | *R* | *T* | *R* | *T* |
| Pollen | 0.60 ± 0.06 | **0.79** ± 0.05 | 0.09 ± 2e<sup>-2</sup> | **0.35** ± 0.04 | 0.28 ± 0.08 | **0.61** ± 0.13 | 0.76 ± 0.0 | **0.95** ± 3e<sup>-16</sup> |
| Darmanis | 0.52 ± 0.07 | **0.72** ± 0.09 | **0.27** ± 0.03 | **0.28** ± 0.05 | 0.24 ± 0.12 | **0.30** ± 0.15 | 0.49 ± 1e<sup>-16</sup> | **0.65** ± 2e<sup>-16</sup> |
| Usoskin | 0.09 ± 0.06 | **0.45** ± 0.16 | 0.10 ± 2e<sup>-2</sup> | **0.12** ± 1e<sup>-2</sup> | 0.07 ± 0.08 | **0.47** ± 0.16 | 0.02 ± 0.0 | **0.69** ± 1e<sup>-16</sup> |
| Mouse Pan. | 0.29 ± 0.05 | **0.56** ± 0.12 | 0.15 ± 0.02 | **0.61** ± 0.03 | 0.37 ± 0.13 | **0.45** ± 0.10 | 0.21 ± 0.0 | **0.48** ± 1e<sup>-16</sup> |
| Muraro | 0.56 ± 0.08 | **0.74** ± 0.12 | 0.40 ± 0.04 | **0.79** ± 0.04 | 0.36 ± 0.15 | **0.44** ± 0.11 | 0.62 ± 1e<sup>-16</sup> | **0.92** ± 0.0 |
| Limb | 0.14 ± 0.03 | **0.66** ± 0.14 | 0.05 ± 0.03 | **0.68** ± 0.12 | 0.10 ± 0.08 | **0.53** ± 0.09 | 0.008 ± 1e<sup>-18</sup> | **0.57** ± 1e<sup>-16</sup> |
| Trachea | 0.11 ± 0.02 | **0.58** ± 0.15 | 0.02 ± 2e<sup>-3</sup> | **0.59** ± 0.0 | 0.05 ± 0.03 | **0.43** ± 0.12 | 0.23 ± 0.0 | **0.45** ± 0.0 |
| Lung | 0.21 ± 0.01 | **0.49** ± 0.09 | 0.12 ± 5e<sup>-2</sup> | **0.71** ± 0.05 | 0.11 ± 0.06 | **0.53** ± 0.10 | 0.19 ± 2e<sup>-17</sup> | **0.42** ± 0.0 |
| Diaphragm | 0.25 ± 0.07 | **0.83** ± 0.17 | 0.09 ± 0.0 | **0.61** ± 0.10 | 0.15 ± 0.11 | **0.63** ± 0.19 | 0.25 ± 0.0 | **0.97** ± 0.0 |
| Spleen | 0.31 ± 0.01 | **0.48** ± 0.08 | 0.21 ± 3e<sup>-2</sup> | **0.28** ± 8e<sup>-2</sup> | **0.41** ± 0.16 | 0.23 ± 0.11 | 0.35 ± 5e<sup>-17</sup> | **0.77** ± 0.0 |

‘R’ and ‘T’ indicate raw and transformed data, respectively. **Bold** indicate higher average accuracy. **Blue** indicate when difference is at most 0.01. Mouse pan. is Mouse Pancreatic dataset.

### p-ClustVal improves clustering with dimension reduction

The empirical evaluation (Table 3) showcases the benefit of *p*-ClustVal in enhancing the performance of widely-adopted dimensionality reduction techniques. For example, using PCA on transformed data leads to maximum improvement of 0.38 on pollen data. With tSNE, maximum gain of 0.36 is achieved on mouse pancreatic data. When using UMAP, maximum gain of 0.62 is seen on spleen data. While the relative performance of dimensionality reduction methods varied across datasets, the empirical analysis consistently underscored the substantial improvements in downstream clustering quality facilitated by the application of the *p*-ClustVal. These findings support the application of *p*-ClustVal as a promising approach for refining dimensionality reduction in the context of high-dimensional genomics data analysis.

**Table 3:**
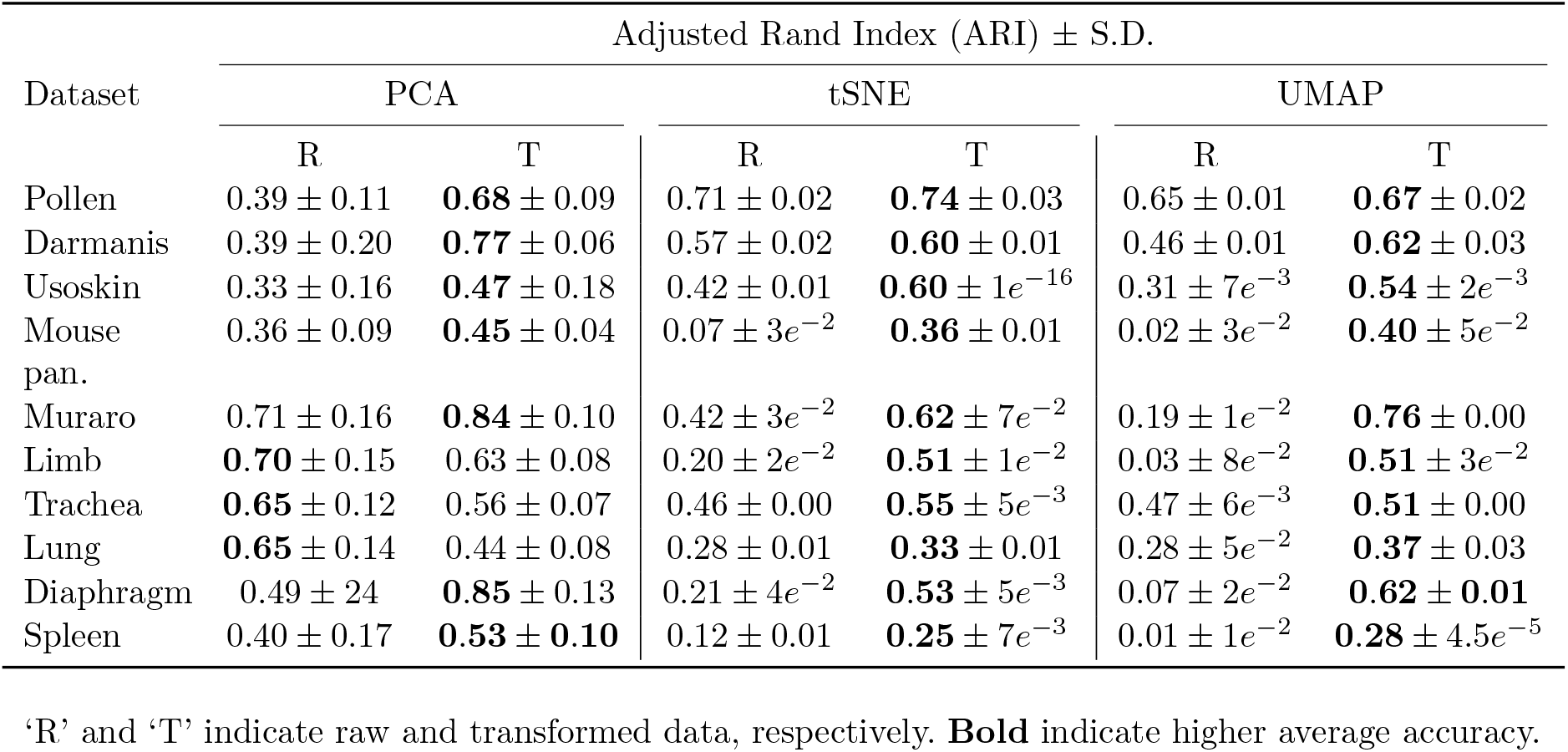
ARI and standard deviation post dimension reduction on raw versus transformed data.

### p-adic-transformation augments SOTA domain specific clustering

Table 4 presents the empirical results, revealing reasonable variation in the accuracy of different tools when analyzing the same dataset, with no single method unanimously outperforming others. However, on most datasets, *p*-ClustVal either significantly improves performance or achieves similar performance as raw data. In Seurat, performance is conserved on 6 out of 10 datasets. RaceID achieves better accuracy on transformed data for 7 out of 10 datasets, with an impressive gain of 0.84 on Diaphragm data, while raw data and Muraro datasets. SIMLR’s transformed data outperforms raw data on all 10 datasets, with a maximum improvement of 0.60 on Lung data. SC3 conserves accuracy on 6 out of 10 datasets, and ADClust’s transformed data performs much better on 9 datasets, with a dramatic accuracy margin of 0.97 points on Diaphragm data. In summary, out of 50 independent observations, using transformed data achieves better or similar results in 86% of cases, while raw data is marginally better in only 14% cases.

**Table 4:** Clustering accuracy of SOTA algorithms on raw and transformed data.

| Dataset | Adjusted Rand Index (ARI) |  |  |  |  |  |  |  |  |  |
| --- | --- | --- | --- | --- | --- | --- | --- | --- | --- | --- |
|  | Seurat |  | RaceID |  | SIMLR |  | SC3 |  | ADClust |  |
|  | R | T | R | T | R | T | R | T | R | T |
| Pollen | 0.77 | **0.8** | 0.58 | **0.82** | 0.09 | **0.5** | **0.97** | **0.97** | 0.10 | **0.68** |
| Darmanis | 0.61 | **0.66** | 0.46 | **0.51** | 0.59 | **0.84** | **0.95** | 0.84 | 0.23 | **0.84** |
| Usoskin | **0.64** | 0.57 | NA | NA | NA | NA | **0.88** | **0.88** | 0.18 | **0.27** |
| Mouse pan. | **0.46** | **0.46** | **0.31** | 0.25 | 0.29 | **0.47** | **0.41** | 0.36 | 0.54 | **0.58** |
| Muraro | **0.93** | **0.93** | **0.83** | 0.78 | NA | NA | **0.94** | **0.93** | 0.39 | **0.93** |
| Limb | **0.66** | **0.67** | 0.5 | **0.56** | 0.24 | **0.52** | 0.64 | **0.94** | 0.04 | **0.58** |
| Trachea | **0.95** | 0.93 | 0.49 | **0.54** | 0.31 | **0.51** | **0.48** | **0.49** | 0.18 | **0.90** |
| Lung | **0.6** | **0.6** | 0.17 | **0.41** | 0.16 | **0.76** | **0.48** | **0.49** | 0.16 | **0.59** |
| Diaphragm | **0.98** | **0.98** | 0.13 | **0.97** | 0.31 | **0.65** | **0.99** | **0.99** | 0.02 | **0.98** |
| Spleen | **0.89** | **0.88** | 0.40 | **0.44** | 0.22 | **0.5** | NA | NA | **0.63** | 0.30 |

‘R’ and ‘T’ indicate raw and transformed data, respectively. **Bold** indicate better results and **blue** indicate either similar performance or when difference is less than 0.01. NA indicate method failed to run on specific data.

## 6 Limitation

Finally, we address the limitations of our approach and discuss the possible solutions. As mentioned in Sec. 1, various factors impact the clustering of scRNASeq data (noise, sparsity, biological and technical variation etc.). However, *p*-ClustVal address the clustering from a data-centric perspective (by preparing an alternate data representation), hence, primary limitation arise due to data-driven parameter estimate. Specifically, *p*-ClustVal is influenced by the accuracy of the `coh` and `sep` parameters, as these parameters directly impact the scaling and relative discretization of data. As described in Sec. 3, `coh` is determined by identifying the neighborhood of each data point. This approach inherently assumes that a data point’s neighborhood primarily consists of data from the same cluster, implying homogeneity. However, this assumption may not hold in datasets with very high variance, where different clusters may frequently overlap, resulting in heterogeneous neighborhoods. This heterogeneity can lead to inaccurate estimates of `coh`, as the top *k* neighbors might include data points from other clusters, skewing the estimate towards higher variance data.

For example, on Adam data with higher overlap between classes, *p*-ClustVal showed marginal improvement over raw data (Table 5). Recently, feature extraction methods, which classify points in an alternate, lower-dimensional space, have been shown to enhance clustering [69, 70]. Consequently, we show that it’s possible to recover the performance of *p*-ClustVal by doing parameters search in a low-dimensional space where neighborhood is relatively homogeneous. It’s important to note that the homogeneity assumption is a reasonable simplification of the problem, allowing for a data-centric, user-independent search for `coh` and `sep` based solely on the data. Moreover, empirical validation has shown that the assumption is effective on majority of datasets considered in this study.

**Table 5:** Adam dataset. *m*: 3660, *d*: 12976, Sparsity: 86.07%

| Dataset | Default |  | PCA |  | tSNE |
| --- | --- | --- | --- | --- | --- |
|  | R | T | T |  | T |
| Adam | 0.02 | 0.09 | 0.21 |  | **0.33** |

‘R’ and ‘T’ indicate raw and transformed data. **Bold** indicate better results.

To better illustrate this, Fig. 7-A visualizes the Adam dataset on the top two principal components, with data colored according to true class labels. The high degree of overlap between different clusters is evident. A corresponding t-SNE visualization (Fig. 7-B) show that clusters are more distinct after t-SNE, making it a better candidate for parameter selection. We conducted a neighbor search in the default data space, PCA and t-SNE space, followed by 20 trials of *k*-means. The results, presented in Table 5, indicate that the default strategy only marginally improves over raw data, whereas parameters derived from PCA and t-SNE based neighbor searches yield reasonable improvement. These finding opens avenues for further research into automated methods that leverage such representations to enhance parameter search and clustering performance.

**Fig. 7:**
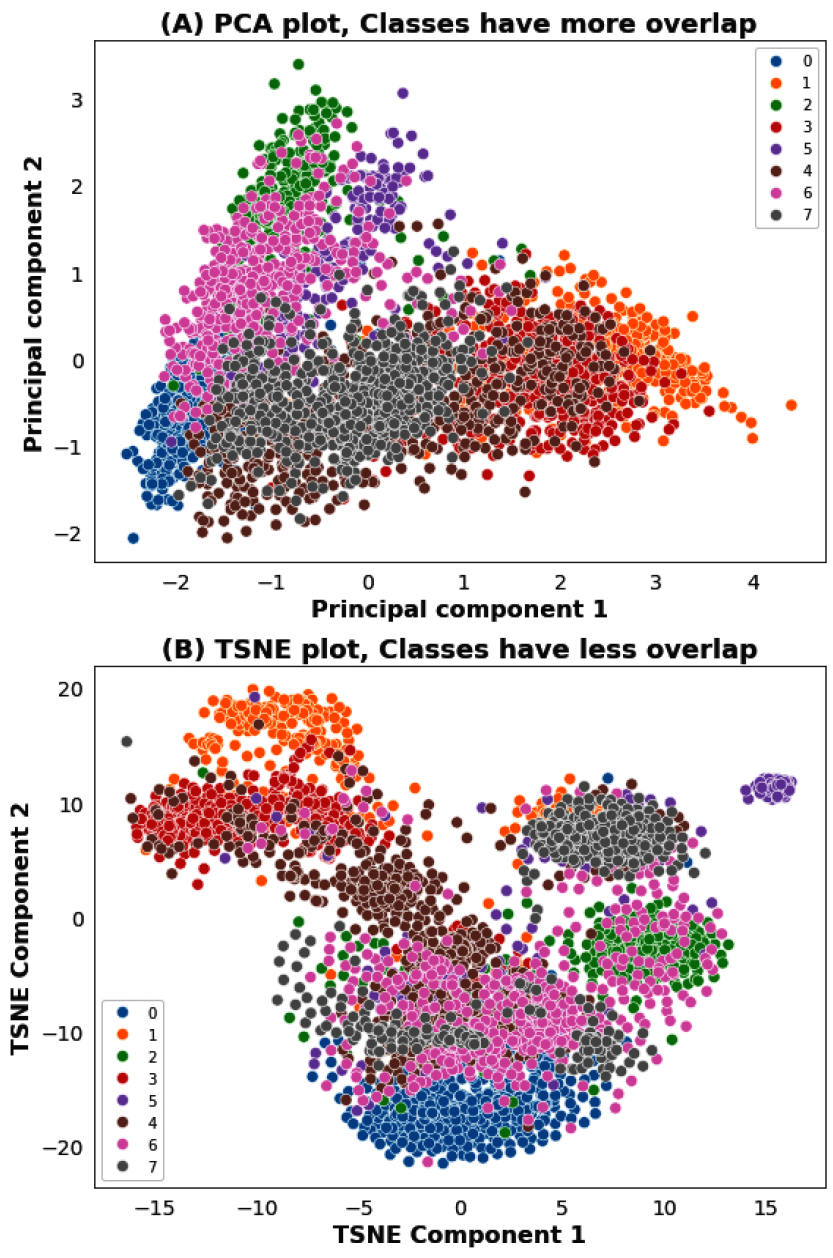
Visualizing the Adam data. Compared to PCA, TSNE reduces the relative overlap of classes.

## 7 Summary & future work

In this work, we introduced *p*-ClustVal, an innovative approach towards creating alternate representations of high dimensional genomics data. Drawing inspiration from the mathematical theory of *p*-adic numbers, *p*-ClustVal leverages metric spaces based on *p*-adic-valuation to enhance cluster separation and discernibility. *p*-ClustVal combined with an unsupervised heuristic for parameter estimation, is validated on downstream clustering, and dimension reduction methods, minimizes tuning requirements and shows remarkable adaptability across various datasets. Comprehensive evaluation on SOTA domain-specific tools demonstrates the robustness of this approach across datasets, and diverse clustering techniques. While *p*-ClustVal currently faces challenges with high cluster overlap, our future direction involves integrating more sophisticated low dimensional embedding to refine data representations, aiming to overcome these limitations. *p*-ClustVal is a novel example of applying abstract mathematical concepts for solving complex real-world problems, and opens up new avenues for interdisciplinary research in mathematics, data science, and biology.

## Acknowledgements

Authors thank the technical support extended by the system administration team of Luddy School of Informatics, Computing & Engineering for their help in deploying the system infrastructure for this study.

## Author Contributions

PS: Conceptualization, Methodology, Development & Design, Experiment, Investigation, Writing – original draft, Writing – review & editing, Visualization. SM: Methodology, Development & Design, Validation, Investigation, Writing – original draft, Writing – review & editing. HK: Methodology, Investigation, Supervision, Writing – original draft, Writing – review & editing. MD: Writing – original draft, Writing – review & editing, Resources, Supervision.

## Code & Data availability

The software code and documentation is available at the Github repository. Relevant data sources are cited in the manuscript.

## Declarations

### Ethics approval

Not applicable.

### Consent to participate

Not applicable.

### Consent for publication

All authors gave their consent for the publication.

## Appendix A Determining number of neighbors (*k*)

Table A1 reports the ARI on different values of *k* (neighbors) for 5 different datasets. The ARI is in similar range despite changes in *k*, and we opted for *k* = 5 in our experiments.

**Table A1:**
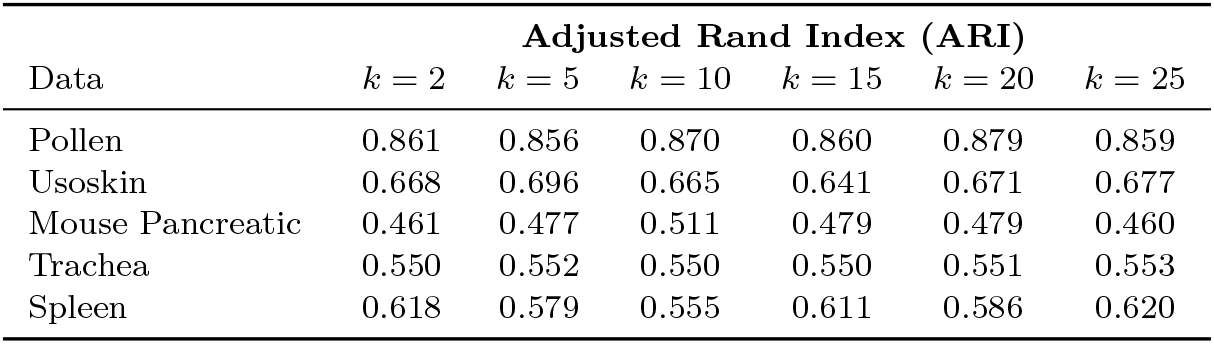
Comparing ARI on different number of neighbors.

## Appendix B Motivation for Data-Centric Parameter Search

The data-driven parameters have been shown to improve methods in learning [71, 72]. [72] argues that data-driven parameters with deep autoencoders and Gaussian process modeling can accurately approximate the wind-turbine wake flow field. [71] has used conditional generative adversarial networks (cGANs) to find solutions to partial differential equations (PDEs) and thus estimate the parameters of the model. They have demonstrated that data-centric parameter estimation can improve accuracy as well as speed. Thus, we also provide a transformation that is tuned with parameters driven by data. The main benefit of data-centric parameter estimation is that it reflects the data more accurately. Compared to the generalized approach where all parameters are one size fits all, this provides more flexibility in tuning.

In our work, we created a transformation function that leverages neighborhood information to infer separation and cohesion parameters. These parameters are then used to discretize the space for the dataset. The approach is unique to each dataset. Contrary to other modeling approaches, which rely on an understanding of the system, data-centric parameter approaches learn directly from the data, providing more freedom. These approaches can be useful in systems such as single-cell RNA-seq, where the underlying system is very complex, making modeling very difficult [73, 74]. Empirically, we have shown improvements in different classes of clustering methods using our transformation.

## Appendix C Data processing

For traditional clustering and dimension reduction, standard data processing was done, for example: genes expressed in less than 1% cells are removed from data followed by removal of cells that have less than 100 genes expressed (non-zero count in less than 100 genes). For data normalization (adjusting for differences in gene counts between cells), we use the method proposed in [75] which ensure that total gene counts in all cells is the same-this removes the bias in cell size (total gene counts of cells) due to sequencing technology or library preparation steps. Finally, *p*-ClustVal is applied on the normalized data.

For experiments with scRNA-seq packages, in Seurat, cells that have non-zero count in more than 100 genes are kept and genes expressed in less than 3 cells are removed. Normalization is done on filtered data by dividing the gene count of each cell by total gene count of that cell and multiplying by the scale factor of 10^4^. The normalized data (*D*) is log transformed as *log*(*D* + 1). For SIMLR and RaceID, data is processed by the default pipeline of the package. SC3 and ADClust does not perform explicit data filtering, so data is processed similar to traditional clustering experiments.

## Appendix D Computing Platform

Experiments on real and synthetic data were done on a 64-bit Ubuntu Linux system with 1.5 TB of memory and 24 dual socket Intel Xeon cores.

